# *In vivo* virulence characterization of pregnancy-associated *Listeria monocytogenes* infections

**DOI:** 10.1101/337444

**Authors:** Holly A. Morrison, David Lowe, Jennifer R. Robbins, Anna I. Bakardjiev

**Affiliations:** Benioff Children’s Hospital, Microbial Pathogenesis and Host Defense Program, University of California, San Francisco, CA 94143; Department of Biology, Xavier University, Cincinnati, OH; Current address: VIR Biotechnology, San Francisco CA, 94158; Current address: 10x Genomics, San Francisco, CA 94105

## Abstract

*Listeria monocytogenes* is a foodborne pathogen that infects the placenta and can cause pregnancy complications. Listeriosis infections usually occur as sporadic infections, but large outbreaks are also reported. Virulence from clinical isolates is rarely analyzed due to the large number of animals required, but this knowledge could help guide the response to an outbreak. We implemented a DNA barcode system using signature tags that allowed us to efficiently assay variations in virulence across a large number of isolates. We tested 77 signature-tagged clones of clinical *L. monocytogenes* strains from 72 infected human placentas and five immunocompromised patients, all isolated since 2000. These strains were tested for virulence in a modified competition assay in comparison to the laboratory strain 10403S. We used two *in vivo* models of listeriosis: the non-pregnant mouse and the pregnant guinea pig. Strains that were frequently found at high abundance within infected organs were considered “hypervirulent,” while strains frequently found at low abundance were considered “hypovirulent.” Virulence split relatively evenly among hypovirulent, hypervirulent, and strains equally virulent to 10403S. The laboratory strain was found to have an intermediate virulence phenotype, supporting its suitability for pathogenesis studies. Further, we found that splenic and placental virulence are closely linked in both guinea pig and mouse models. This suggests that outbreak and sporadic pregnancy-associated *L. monocytogenes* are not generally more virulent than lab reference strains. However, some strains did show consistent and reproducible virulence differences, suggesting that their further study may reveal deeper insights into the biological underpinnings of listeriosis.

## Introduction

Listeriosis is a foodborne disease that afflicts humans worldwide (1, 2). In the United States, the Centers for Disease Control estimates it is responsible for approximately 1,600 cases and 260 deaths per year (3). Most cases occur in predisposed individuals such as immunocompromised patients, neonates and elderly adults. In those cases the main clinical manifestations are sepsis, meningoencephalitis, and death (4). With a mortality rate of ∼20% and recurring foodborne outbreaks, listeriosis remains a significant public health concern (2, 5–7).

Disseminated infections are of particular concern in pregnant women, as *Listeria monocytogenes* can spread to the placenta, fetus and/or neonate. Approximately 14% of clinically recognized cases occur during pregnancy (8). Infection may lead to pregnancy loss, preterm birth, stillbirth, and life-threatening neonatal infections (9); however, the mechanisms by which *L. monocytogenes* reaches and breaches the placenta are only just beginning to be understood using animal models (10). We previously established the pregnant guinea pig model of listeriosis, which mimics human disease (11). After intravenous inoculation, the maternal spleen and liver are colonized rapidly, whereas the placenta greatly resists *L. monocytogenes* infection and is delayed in colonization (12, 13). It is possible that the placenta can only be infected after robust dissemination of the bacteria throughout maternal organs. Alternatively, or additionally, it is possible that pregnancy-associated cases of *L. monocytogenes* represent bacterial strains that are more virulent generally or more specifically adapted for placental colonization.

*L. monocytogenes* typically has a saprophytic lifestyle and is commonly found in soil, vegetation, and animal feces. Furthermore, it is highly resistant to common antibacterial precautions taken in food preparation; e.g. cold temperatures, desiccation, and high salt. These factors combine to make *L. monocytogenes* a common food pathogen, but the infectious dose is high, and so most cases of listeriosis are isolated, sporadic events (8). Indeed, the average adult ingests ∼10^5^ CFU four times a year, but only a small number of predisposed individuals contract listeriosis (14). Occasionally, major outbreaks occur in widely distributed foods, leading to larger numbers of infections (5, 6). It remains an open question whether these outbreak strains are more virulent than sporadic or lab reference strains.

Increasingly, we are learning about how outbreak and hypervirulent pathogen strains arise and diverge from reference lab strains through the burgeoning field of microbial population biology. Several studies have analyzed pathogenic strains to understand their evolution and population structure (15–19), and some assay the virulence of representative clonal clusters relative to historical reference strains (20). While these studies identify molecular differences between strains that can account for their origin and altered virulence, actually assaying their virulence *in vivo* is challenging due to the large number of laboratory animals required. This is especially true when considering the testing of clinical isolates, with strains numbering in the scores or hundreds. However, the use of DNA barcodes (signature tags) can allow for multiplexed analysis of several strains within a single animal. Such studies allow researchers to understand how virulence has evolved in clinical isolates over time while comparing them to lab reference strains.

Here we characterize the virulence of 77 *L. monocytogenes* strains: 73 isolated from sporadic clinical cases over a 10-year period and four strains isolated from pregnant women infected during outbreaks. Sixty-eight sporadic isolates were from pregnancy-associated listeriosis cases, while five were from non-pregnant immunocompromised patients. We set out to identify strains with increased and decreased systemic virulence as compared to lab references, using a barcode-based competition assay in pregnant and non-pregnant animal models. We also assayed for trends in virulence, comparing bacterial burdens across organs to determine which maternal organs were most likely to be infected in concert with the placenta.

## Materials and Methods

### Bacterial strains and culture conditions

The laboratory reference strains are 10403S (erythromycin susceptible) (21), DP-L3903 (erythromycin resistant) (22), and signature-tagged 10403S strains (23). All *L. monocytogenes* clinical strains used in this study are listed in Supplementary Table S1. Sixty-eight clinical isolates of *L. monocytogenes* from pregnancy-associated listeriosis that occurred over 10 years (2000-2010) in 25 states in the US were obtained from the Centers for Disease Control and Prevention (CDC, Atlanta, GA). Five strains isolated from the blood of immunocompromised patients at Memorial Sloan-Kettering Cancer Center were a generous gift from Dr. Michael Glickman. Bacteria were grown in brain heart infusion (BHI, Bacto®, BD) media at 37°C. When necessary, media were supplemented with the following antibiotics, all purchased from Sigma: chloramphenicol (7.5μg/mL), nalidixic acid (25μg/mL), streptomycin (200μg/mL) or erythromycin (2μg/mL).

### Signature tag (DNA barcode) integration in clinical strains

Unique 40-bp signature tags (STs) were inserted into *L. monocytogenes* strain genomes by site-specific integration from the pPL2 vector as previously described (23). Tagged clinical strains generated in this study used tags 116, 119, 191, 205, 210, 219, 231, 234, 242, 288 and 296. Integrations were confirmed by selection for chloramphenicol resistance and PCR as previously described (24).

### Animal infections

This study was carried out in strict accordance with the recommendations in the Guide for the Care and Use of Laboratory Animals of the National Institutes of Health. All protocols were reviewed and approved by the Animal Care and Use Committee at the University of California, San Francisco (IACUC# AN079731-03A). Individual strains were grown in BHI at 37°C overnight. On the day of infection, 11 differentially-tagged strains were combined at equal ratios to generate ten input pools. Nine input pools (clinical pools) contained nine clinical and two 10403S strains; one input pool (control pool) contained 11 differentially-tagged 10403S strains. 6-8 week old non-pregnant female CD1 mice (Charles River Laboratories) were inoculated i.v. with a total of 2×10^5^ CFU pooled bacteria per animal. Pregnant Hartley guinea pigs (Elm Hill Labs, MA) were inoculated i.v. on gestational day 35 with a total of 1×10^8^ CFU pooled bacteria per animal. For the mouse experiments, each clinical pool was injected into five mice on two separate days for a total of ten mice per pool; the control pool was injected into 15 mice on three separate days. Murine spleens were removed at 48 h.p.i. For the guinea pig experiments each pool was injected into 2-5 pregnant guinea pigs depending on the number of fetuses per dam. The total number of guinea pigs injected with clinical pools was 24 with a total of 96 placentas. The control pool was injected into 3 guinea pigs with a total of 11 placentas. Guinea pig spleens and placentas were removed at 24 h.p.i. Organs were homogenized in 0.2% Igepal (Sigma) with a tissue grinder. Aliquots from each output pool were plated on BHI agar plates containing 25μg/mL nalidixic acid. CFU per organ were enumerated, and at least 10^4^ colonies from each output pool were scraped off the plates and re-suspended in PBS. Aliquots of these suspensions were stored at −20°C. Input pools were prepared in the same fashion.

### qPCR

Genomic DNA was extracted from input and output pools using a Gram-positive DNA purification kit (Epicentre), substituting mutanolysin (5U/μL, Sigma) for lysozyme. Relative quantification by qPCR for each signature tag was achieved with previously published primer sets: signature tag-specific forward primers and the common pPL2-395R reverse primer (23). In addition, one primer set (LIM2 and LIMRE) was directed against *iap*, a gene used as internal reference (25). All qPCR reactions were performed in a Roche LightCycler® 480 qPCR machine. Each 20μL reaction contained 10μL SsoAdvanced™ SYBR® Green Universal Supermix (Bio-Rad), 200nM of each primer, nuclease-free water and template DNA. A total of 20ng template DNA was used for experimental samples. DNA extracted from 10403S-signature tagged reference strains was used to construct qPCR standard curves for each signature tag primer set, with template amounts of 100ng, 10ng, 1ng, 0.1ng, and 0.01ng. Cycling conditions were as follows: 98°C 2’, (98°C 5’’, 60°C 20’’, 68°C 20’’) x 40 cycles, followed by a melting curve cycle (98°C 15’’, 60°C 30’’, ramp to 98°C in 0.29°C/sec intervals). For each animal species, duplicate qPCR reactions for the standard curve dilutions, input and output pools, and template-free controls were run in parallel on a single 384-well plate per primer set.

The relative abundance of each signature tag in each output sample was determined in relation to the reference gene *iap* and the respective input pool. Quantification of cycle numbers and primer efficiencies were obtained using Lightcycler® Software release 1.5.0 SP3 (Roche). Relative abundance (RA) values were calculated using the following equation, which accounts for different primer efficiencies (26): RA = ((E_iap_^Cq_iap-sample_)/(E_ST_^Cq_ST-sample_))/((E_iap_^Cq_iap-input_)/(E_ST_^Cq_ST-input_)), where E_iap_ and E_ST_ are the efficiency values calculated from the standard curves for the *iap* and ST-specific primers.

### Determination of virulence

Within each output pool, the average relative abundance was calculated for each clinical strain and divided by the average abundance of the two reference strains in the same output pool. This yielded an output pool-specific, normalized relative abundance for each clinical isolate. The standard deviation of normalized abundances was calculated using the control group that consisted of 11 differentially-tagged 10403S strains. A z-score describing the normalized relative abundance for each strain compared to 10403S was then calculated by subtracting the mean of the control group relative abundance and dividing by the standard deviation of control group relative abundance. Strains that were significantly more or less abundant (p < 0.01) were identified according to a normal distribution of z-scores.

### Direct competition assay

6-8 week old female CD1 mice (Charles River) were inoculated i.v. with 2×10^5^ CFU of one clinical isolate (erythromycin-susceptible) and 10403S (erythromycin-resistant) at 1:1 ratio. Bacteria were recovered from spleen at 48 h.p.i. and enumerated, then individual colonies were tested for differential susceptibility to erythromycin to represent clinical versus 10403S reference strain. The control group was injected with a 1:1 ratio of two 10403S strains that differed in their susceptibility to erythromycin. Statistical significance was determined by one-way ANOVA with Dunnett’s multiple comparisons post-test.

## Results

### Clinical isolates and in vivo screening method

Our laboratory reference strain 10403S (21) is a streptomycin resistant derivative of *L. monocytogenes* strain 10403, which was originally isolated from a human skin lesion in 1968 (27). 10403S is one of the most widely used strains for experimental investigation and has been passaged for decades under laboratory conditions (28). We sought to use a DNA strain barcoding and pooling assay scheme (Fig. 1) to determine how dozens of recent clinical isolates that had not been previously cultivated in the laboratory differ in virulence from 10403S.

**Fig. 1.**
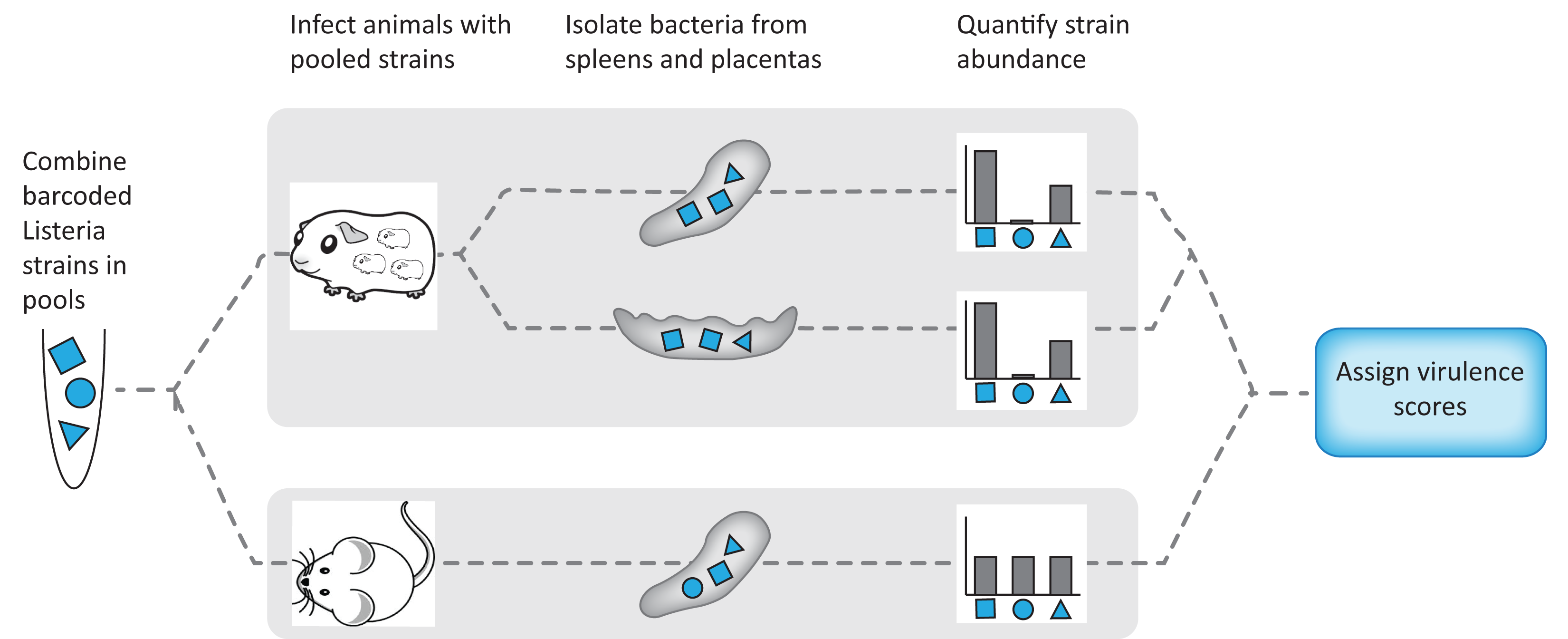
Experimental design. Signature-tagged *L. monocytogenes* strains were pooled and injected i.v. into pregnant guinea pigs or non-pregnant mice. Each pool contained 11 barcoded strains; 9 clinical and 2 laboratory reference strains (10403S) in the clinical pools, and 11 laboratory reference strains in the 10403S pool. For each organ set (guinea pig spleen, guinea pig placenta, mouse spleen), virulence scores were assigned to each strain based on the average relative abundance in the infected organs in comparison to the laboratory reference strains.

We compiled 77 clinical isolates of *L. monocytogenes*: 72 strains from pregnancy-associated cases of listeriosis collected by the CDC over a 10-year period (2001 to 2011) in 24 US states, and five strains from the blood of immunocompromised non-pregnant patients undergoing cancer therapy at Memorial Sloan Kettering Cancer Center (MSKCC) in New York (Fig. 2A and Supplementary Table S1). Almost all strains were from sporadic cases of listeriosis. Four strains were from three different outbreaks of listeriosis associated with the following contaminated food sources: (i) Mexican-style cheese in 2005 (placental isolate, serotype 4b) (29), (ii) turkey deli meat in 2006 (placental and neonatal blood isolates from unrelated mother and neonate, serotype 4b) (30), and (iii) hog head cheese in 2011 (maternal blood isolate, serotype 1/2a) (7). Only the strains from the CDC were serotyped. Among these, serotype 4b was most common, followed by 1/2a and 1/2b, consistent with previous reports (5, 6) (Fig. 2B).

**Fig. 2.**
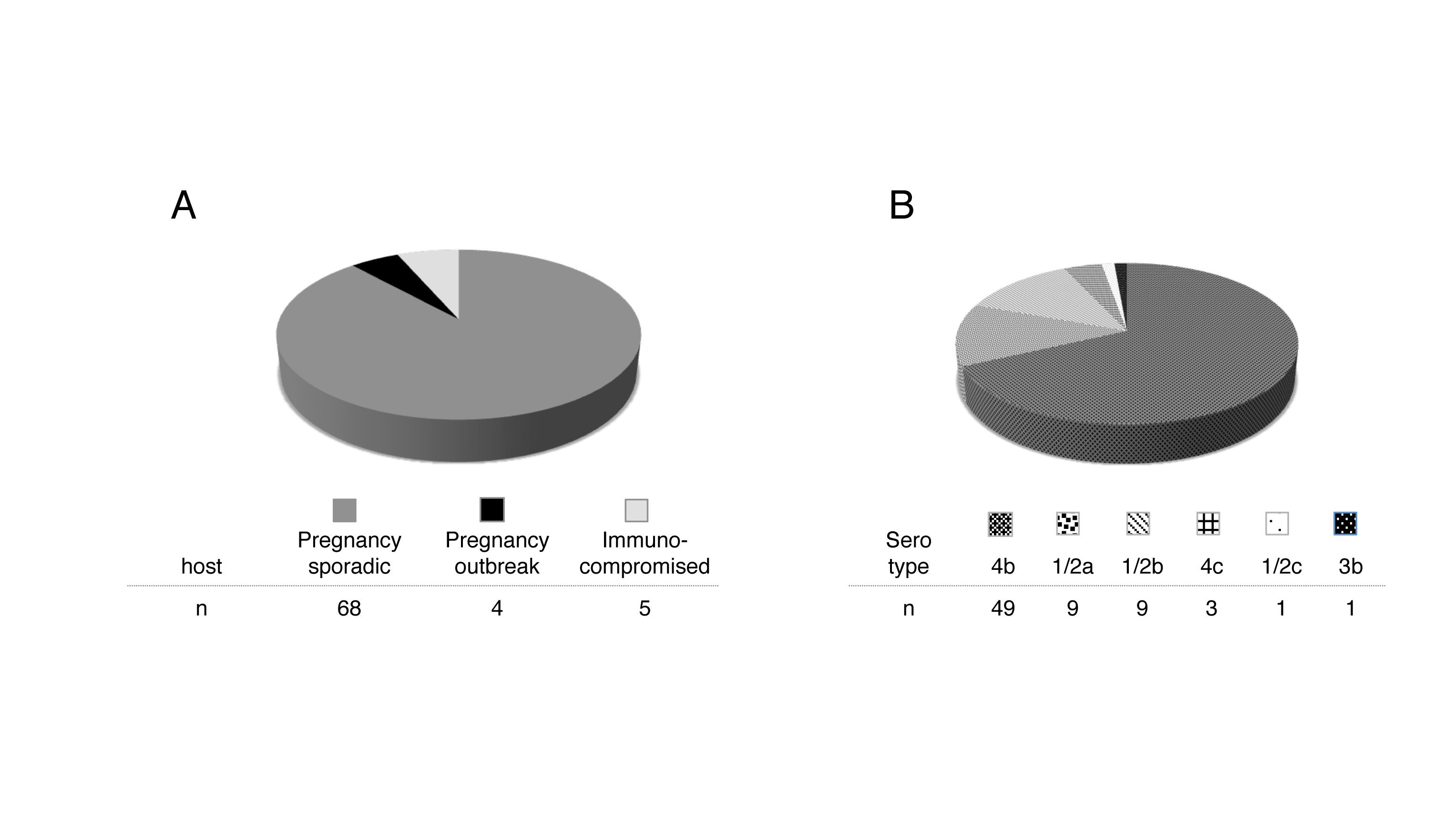
Clinical isolates. **A) Pregnancy associated** *L. monocytogenes* strains (n=72) from 25 US states were collected by the CDC between 2000-2010. Most were associated with sporadic cases of listeriosis during pregnancy and were isolated from placental tissue (n=68; pregnancy, sporadic). Four strains were associated with listeriosis outbreaks in the US (n=4; pregnancy, outbreak). These 4 strains were isolated from placenta (n=2), maternal blood (n=1), and neonatal blood (n=1). Five strains were isolated from immunocompromised patients at MSKCC (n=5; immunocompromised). **B)** Serotype distribution of pregnancy-associated strains.

We compared the virulence of each clinical strain to 10403S in two animal models: (1) non-pregnant mice, the standard model for the pathogenesis of systemic listeriosis; and, (2) pregnant guinea pigs, an excellent small animal model for pregnancy-associated listeriosis (11). In order to minimize the number of animals required for virulence screening, we incorporated a different, previously characterized DNA barcode into the chromosome of each clinical isolate (23). Clinical strains were assigned to pools a priori; pools were balanced such that they included one of each signature tag from the set used, and each included one commonly tagged and one differentially tagged 10403S strain. Subsequently, each animal was inoculated with pools of differentially-tagged bacteria. We used a total of 10 pools, each containing 11 strains marked by unique barcodes. The control pool contained eleven 10403S strains, while each of the remaining nine pools consisted of nine clinical and two 10403S strains (Pools A-I).

### Profiling systemic virulence in mice and guinea pigs

Mice were infected intravenously (i.v.) with a total of 2×10^5^ CFU/animal (10 animals/pool). The median bacterial burden in the control spleens 48 hours post-inoculation (h.p.i) was 7.2×10^7^ CFU (Fig. 3A). The median CFU in the spleen of mice inoculated with pools containing clinical strains ranged from 5.6×10^7^ CFU (Pool D) to 1.9×10^8^ CFU (Pool G), and did not differ significantly from the median of the control pool except in two instances: the median bacterial burden of Pools F and G were 1.8- and 2.6-fold higher than the control pool.

**Fig. 3.**
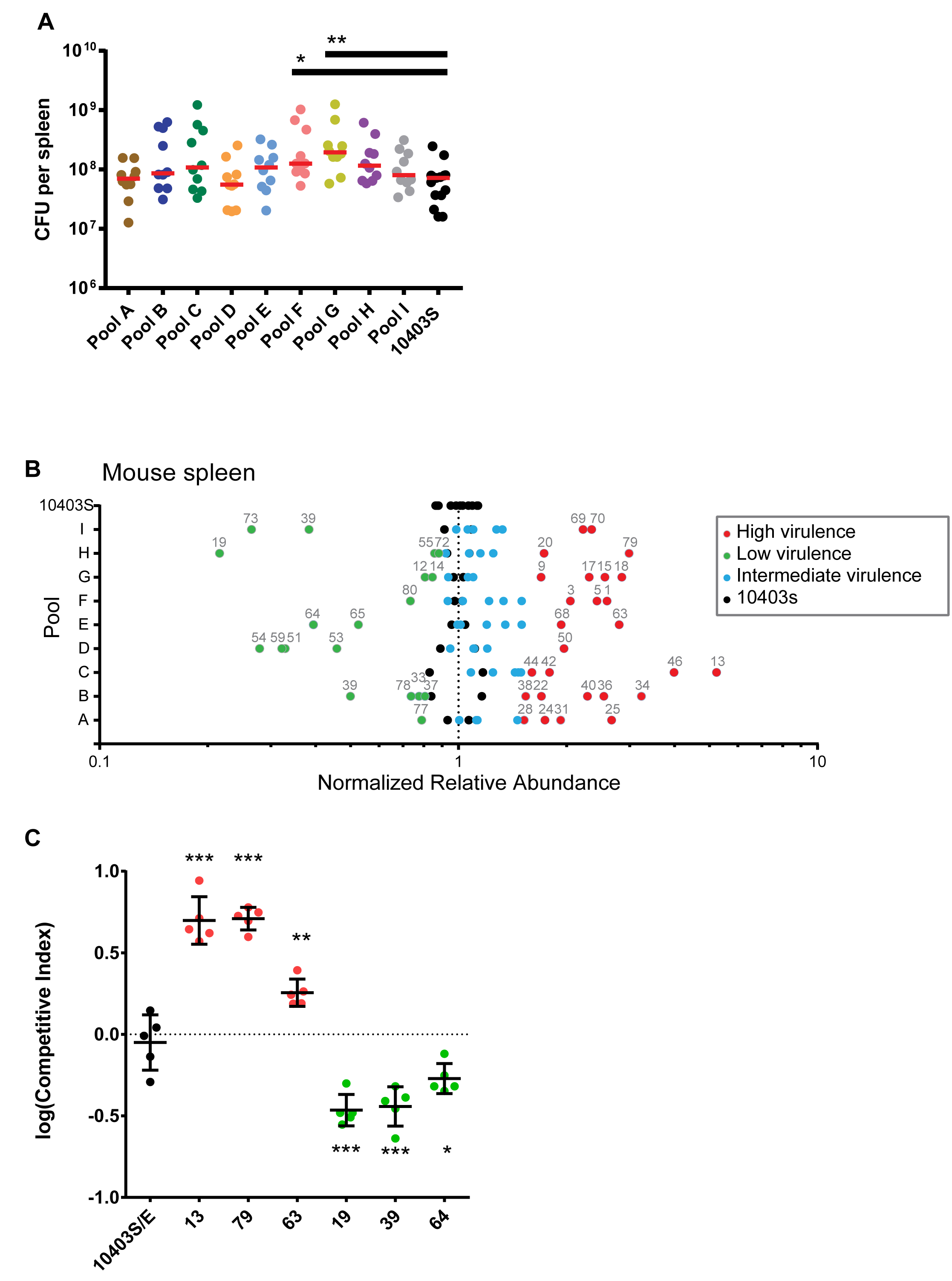
Virulence screen of clinical *L. monocytogenes* isolates in murine spleen. CD1 mice (non-pregnant) were infected i.v. with bacterial pools containing differentially-tagged *L. monocytogenes* strains at equal ratios (total of 10 pools). Pools A-I contained nine clinical and two 10403S strains per pool; the 10403S pool contained 11 laboratory reference strains. Statistically significant differences in splenic bacterial burden compared to the control group were determined using one-way ANOVA with Dunnett’s multiple comparisons post test. ***, p<0.0001. **, p<0.01. *, p<0.05. **A)** Bacterial burden in murine spleen 48 h.p.i. with 2×10^5^ CFU per pool. Pools A-I: n=10 mice/pool; 10403S pool: n=15 mice. Each circle represents the bacterial burden in one spleen, and each pool is represented by a different color. Red lines represent median. **B)** The average relative abundance of each strain in mouse spleen was quantified by qPCR. To accurately compare values across pools, the average relative abundance for each isolate was then normalized to the average of the reference strain in each pool. Significance z-scores were calculated for the deviation from the range expected based on the 10403S pool (black circles). Blue circles indicate isolates with virulence similar to 10403S (intermediate virulence). Red and green circles indicate isolates with significantly higher and lower virulence, respectively. **C)** CD1 mice were infected with one erythromycin-resistant 10403S strain and one erythromycin-susceptible clinical isolate at a 1:1 ratio. The clinical isolates were chosen based on their virulence scores in Fig 3B: 3 hyper-(red circles) and 3 hypo-(green circles) virulent strains. Competitive indices (isolate/10403S) were calculated for bacteria recovered from the spleen 48 h.p.i. The control group was infected with two 10403S strains that differed in their susceptibility to erythromycin (10403S/E, black circles). Each group contained 5 mice from 2 separate experiments.

Using qPCR with primers specific for each DNA barcode, we determined the average relative abundance of each clinical strain in comparison to 10403S among the bacteria recovered from each spleen (Fig. 3B). We observed a range of virulence phenotypes both within and across the individually analyzed pools. We found that 27 strains were significantly more virulent (z-score >2.0, red points in Fig. 3B) and 18 strains were significantly less virulent (z-score <-2.0, green points in Fig. 3B) than 10403S. Strains with significantly different virulence were present in all pools. Most pools contained one or more high and low virulence strains; only one pool did not contain a low virulence strain (Pool C). Importantly, four sporadic clinical strains (strains 2, 16, 21, and 39; see also Supplementary Table S1) that were present in two different pools showed similar virulence in their two pools, suggesting that the combination of strains within each pool did not significantly influence the virulence score of individual strains.

We validated our approach by direct competition of select clinical isolates with 10403S in non-pregnant mice (22). We chose six clinical strains with virulence scores that were either significantly higher or lower than 10403S in the pooled assay. Mice were inoculated i.v. with one clinical isolate in combination with 10403S, and their spleens assayed for bacteria at 48 h.p.i. The strains differed in their susceptibility to erythromycin and were injected at a ratio of 1:1 and a total CFU of 2×10^5^/mouse. Consistent with the results of our screen, the two hypervirulent strains 13 and 79 were ∼5-fold more virulent than 10430S and strain 63 was 2-fold more virulent (Fig. 3C). In contrast, the hypovirulent strains 19, 39, and 64 were 2-3-fold less virulent than 10403S. These results recapitulated the virulence phenotypes identified in the screen.

Next, we infected pregnant Hartley guinea pigs i.v. with 1×10^8^ CFU of the same pools we used in the mouse screen, and determined the bacterial burden 24 h.p.i. We chose an earlier time point than in the mouse screen to avoid the potentially confounding effect of bacterial trafficking between placenta and spleen at later time points (12). Twenty-four pregnant guinea pigs were inoculated with clinical pools (2-5 animals/pool); 3 animals were inoculated with the control pool. The median bacterial burden in the spleens of the control pool was 2.4×10^6^ CFU, and ranged from 3.6×10^6^ CFU (Pool D) to 3.1×10^7^ CFU (Pool C) in the spleens of animals inoculated with pools containing clinical isolates, indicating higher overall burdens (Fig. 4A). We determined the average relative abundance of each strain in the guinea pig spleen normalized to 10403S as described above. We identified 22 hypervirulent and 20 hypovirulent strains (Fig. 4B).

**Fig. 4.**
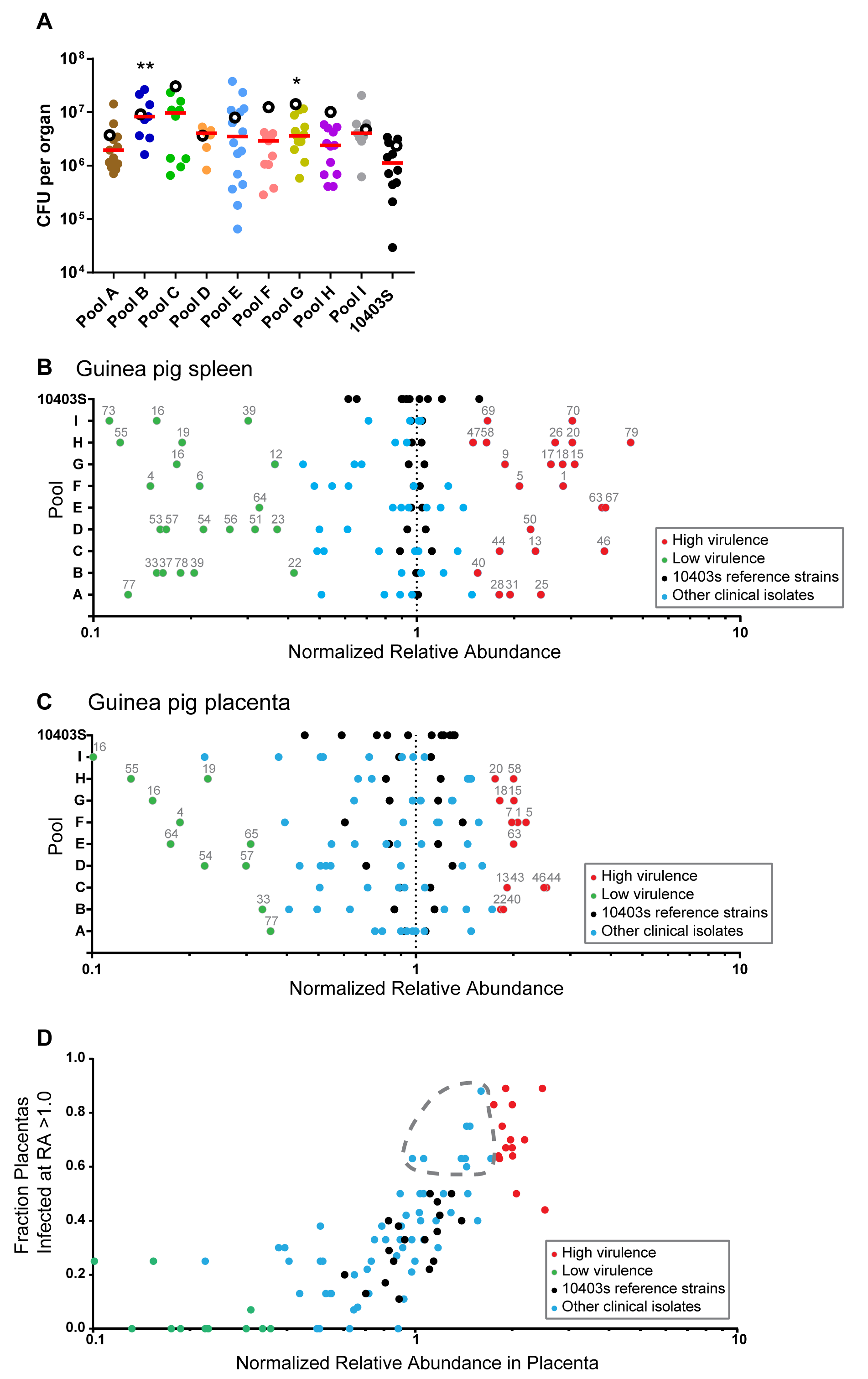
Virulence screen of clinical *L. monocytogenes* isolates in pregnant guinea pigs (spleen and placenta). Pregnant Hartley guinea pigs were infected i.v. with pools containing differentially-tagged *L. monocytogenes* strains (see Fig. 3). Statistically significant differences compared in bacterial burden in spleen and placenta to the control group were determined using one-way ANOVA with Dunnett’s multiple comparisons post-test. ***, p<0.0001. **, p<0.01. *, p<0.05. **A)** Bacterial burden in guinea pig spleen and placenta 24 h.p.i. with 10^8^ CFU per pool. The total number of guinea pigs was 27 with a total of 107 placentas. Number of placentas in each pool: A=12; B=8; C=9; D=8; E=15; F=10; G=14; H=12; I=8, 10403S=11. Each filled circle represents the bacterial burden in one placenta, and each pool is represented by a different color. Red lines represent median placental CFU. Empty circles represent the median bacterial burden in spleens from each pool. **B)** The average relative abundance of each strain in guinea pig spleen was quantified by qPCR and significance z-scores were calculated. Black dots indicate 10403S strains. Blue circles indicate isolates with virulence similar to 10403S (intermediate virulence). Red and green circles indicate isolates with significantly higher and lower virulence, respectively. **C)** Average relative abundance of each strain in guinea pig placenta quantified and calculated as described above. **D)** Correlation of relative abundance of each strain in the placenta with the fraction of placentas they infected at higher relative abundance than their inoculant (RA>1.0). Gray dashed outline encircles isolates not identified as highly virulent by relative abundance alone, but with infected fractions comparable to high virulence isolates. Color coding corresponds to panel C.

In both animal models, high- and low-virulence strains were distributed stochastically across the pools, which we expected with randomized pool assignments. In the guinea pig spleen the relative abundance of 10403S in the control pool exhibited a wider range than in the mouse (compare Fig. 3B to 4B). However, the virulence scores of the clinical isolates were similar between mouse and guinea pig spleen. The scores were concordant for 70% (54/77) of the strains, and among the discordant strains all but one were either hyper- or hypovirulent in one animal model and intermediately virulent in the other animal model (Supplementary Table S2). Only one strain (strain 22, an outbreak strain) was hypervirulent in murine spleen and hypovirulent in guinea pig spleen.

### Virulence screen in the guinea pig placenta

We evaluated the relative virulence of the clinical isolates in the placentas (n=107) of the inoculated guinea pigs (8-15 placentas/pool). The median bacterial burden in the control group was 8.2×10^5^ CFU per placenta (Fig. 4A). The median of the clinical pools ranged from 1.7×10^6^ CFU (Pool A) to 8.4×10^6^ CFU per placenta (Pool C). The range of CFU across all placentas spanned 3-log (3×10^4^ to 3.8×10^7^ CFU), which is typical for placental infection and likely due to the stringent bottleneck in placental colonization (12). Consistent with a tight bottleneck we found the bacterial founding population in the placenta to be significantly smaller than in the spleen. We calculated a median founding population of 1.1×10^5^ CFU in spleens and 278 CFU in placentas, respectively (Supplementary Figure S1).

Next, we determined the relative abundance of clinical isolates in the guinea pig placenta in comparison to 10403S. We identified 14 clinical strains with high and 10 clinical strains with low virulence in the placenta (Fig. 4C). As in the spleen, high and low virulence strains were distributed stochastically across the pools. Virulence was also assayed by comparing the fraction of placentas where a strain had a high relative abundance (RA >1) compared to its relative abundance in guinea pig placentas. We reasoned that hypervirulent strains would be able to infect more placentas as well as have greater abundance within placentas. In general, the fraction of infected placentas did correlate strongly with the average relative abundance across placentas (Fig. 4D). However, this analysis also revealed nine strains with a fraction of infected placentas equivalent to or higher than that of several strains deemed more virulent by the relative abundance parameter described above.

Comparison of the virulence scores in placenta and/or spleen of both rodents showed a striking degree of overlap among the three datasets. Only two strains showed a placenta-specific virulence phenotype (strains 7 and 43). These were hypervirulent in the placenta (by Z-score and fraction of infected placentas), and intermediately virulent in spleen of guinea pigs and mice. The five strains that were isolated from immunocompromised, non-pregnant adults all had intermediate virulence scores in the placenta, and varying virulence scores in the spleen of both animal models (Supplementary Table S1). The four outbreak strains demonstrated variable virulence scores across all organs; only one of the outbreak strains scored hypervirulent in all organs. However, due to the small number of these strains it is not possible to draw any further conclusions.

### Discussion

Here we report the *in vivo* virulence phenotypes for 73 sporadic and four outbreak clinical strains of *L. monocytogenes*, 72 of which were isolated from cases of pregnancy-associated listeriosis. Using a novel DNA barcode approach with qPCR, we identified isolates with either significantly higher or lower virulence than the standard laboratory reference strain 10403S in systemic listeriosis as well as placental infection. However, no strain showed more than a 5-fold difference in virulence compared to 10403S. By using signature tagged (barcoded) strains and qPCR, we found the 77 strains to be an even mix of hypervirulent, hypovirulent and intermediately virulent. Both outbreak and sporadic clinical isolates were compared, but neither associated with any virulence phenotype.

Our isolates included four strains collected during recent outbreaks of foodborne listeriosis in the United States (7, 29, 30). In contrast to the bloodstream isolates from septicemic patients, these isolates were each associated with otherwise healthy pregnancies. We observed that one of these strains was highly virulent in all three assays, while the remaining three showed varied but overall moderate virulence patterns (Supplemental Table 1, Strains 13, 21, 22, 23). It is tempting to assume that outbreaks are due to increases in virulence. However, in addition to the bacterial virulence, independent factors such as ingested dose, maternal genetics and overall maternal health may dramatically influence the outcome of exposure to *L. monocytogenes*. Evaluating the effect of any of these factors would require additional studies, potentially including prospective studies to fully characterize the maternal status correlated with placental infection and pregnancy outcomes.

Population biology studies of pathogens have focused primarily on how virulence evolved, outbreaks arose, and antibiotic resistance spread (15–18, 31). Fewer studies have sought to compare the *in vivo* virulence of clinical strains over a period of time. In part, this is due to the high cost of animal research and the need for several animals per strain. In order to circumvent this, we developed a DNA barcode system. Previous uses of signature tagged strains in *L. monocytogenes* have involved understanding bottlenecks in disseminations and alanine suppression screening to investigate virulence factors (13, 23). Here, it allowed for the simultaneous use of clinical strains in order to reduce the number of animals required to assess virulence. This technique could be even more valuable in larger, more expensive animal models, such as nonhuman primates. Additionally, the ability to test resistance to food processing techniques could be streamlined by using signature tagged libraries of clinical strains.

We observed a larger variation in the distribution of strain abundances in the guinea pig placenta than in either of the spleen datasets. This is consistent with the previously reported bottleneck for placental infection (12, 13); therefore, we determined the founding population in the guinea pig placenta. We calculated approximately 1/360,000 bacteria from the inoculum will infect the placenta. Many of the hypervirulent strains both had a higher abundance in the placenta and infected a greater fraction of placentas. Therefore, in assessing virulence for organs in which an infection bottleneck exists, CFU burden alone are an incomplete measure, and the fraction of organs infected should also be evaluated.

Clinical strains had similar virulence between their spleens and placentas. *L. monocytogenes* strains have been analyzed by multilocus strain typing and organized into clonal clusters (18). The most prevalent clonal clusters in bacteremia were also present in placental and neuroinvasive strains. This suggests that successful placental colonization requires a robust systemic infection. It does not mean, however, that *L. monocytogenes* has not evolved specialized determinants to infect the placenta. Guinea pig models have identified genes required for successful colonization of the placenta compared to the liver (32). And outbreak strains in some pathogens have been traced to novel virulence factors through recombination or horizontal gene transfer (33). A notable example is the EHEC O157:H7 strain that gained shiga toxin genes via horizontal gene transfer (34). Further, *Streptococcus* species have novel virulence factors associated with accessory regions; that is, genes not found in the core genome (35). However, *L. monocytogenes* has been reported to have a highly conserved and syntenic genome (36). Out of the large number of clonal clusters, only the CC4 strains have so far demonstrated an increase in neuronal and placental infection without an increase in splenic or hepatic infection, likely due to a novel carbon metabolism operon (20). We only observed one instance of a decreased splenic virulence and increased placental virulence. Interestingly, this strain, LS22, was isolated from neonatal blood during a deli meat outbreak (30). However, another isolate from the same outbreak but isolated from a placenta (LS23) did not show this phenotype. Both strains were serotype 4b, which is more commonly associated with clinical cases (37).

Our lack of strains with increased placental virulence compared to maternal organs may be due to our sample size of clinical isolates being ∼1/100^th^ of that initially used by Maury et al., (20); although that work assayed a similar number of clones for virulence, they were chosen as representative of the starting population’s clonal clusters. The tight linkage between maternal and placental virulence and the fact that human placental infection provides no epidemic selective advantage suggests that placenta-specific strains are likely rare.

Our survey of virulence in both sporadic and outbreak strains from pregnancy-associated listeriosis cases shows that American *L. monocytogenes* isolates are evenly spread around the long-used laboratory strain 10403S, with some more and some less virulent in animal models. This validates the use of that laboratory strain in pathogenesis studies. Further, the lack of clear difference between outbreak and sporadic strains suggest that listerial epidemiology is not a function of pathogen virulence but of other factors, likely related to individual behaviors/health and food production practices. Finally, we found a tight coupling between maternal bacterial burden and placental infection, suggesting that a primary driver of placental susceptibility is the degree of maternal infection. The DNA barcode approach is a powerful and cost-efficient way to assess the performance of large numbers of diverse clones in animal models.

## Acknowledgements

We are grateful to Lewis Graves (CDC) who provided the *L. monocytogenes* strains. This work was supported by NIH R01AI084928 (A.I.B), Burroughs Wellcome Fund (A.I.B), NIH F32AI102491 (H.A.M), and NIH F32AI120676 (D.E.L).

## Supplementary Information

**Table S1. Strains used in this study.**

^1^Strain ID designated by outside laboratory

^1^Date received at CDC

^3^State where isolate was originally acquired

^4^As determined at CDC

^5^strain number used in this study

^6^Shading indicates significant z-scores. Red, z-score >2.0. Green, z-score <-2.0

n/a, not applicable

nd, not determined

**Table S2. Comparison of splenic virulence scores between mouse and guinea pig.**

## Supplementary Figure S1

**Fig. S1. Quantification of the founding population in spleens and placentas of guinea pigs.**

Pregnant Hartley guinea pigs (spleens n = 3; placentas n = 11) were infected i.v. with pools containing differentially-tagged in the same 10403S strain. The founding population (Nb) was calculated by the harmonic mean of the tag abundance based in the organ. A Mann-Whitney test of the the founding populations in spleens and placentas found a statistically significant difference (\*\**P*-value = 0.0055). Filled circles represent the amount of bacteria that founded the infection of the organ in CFU/mL and red bars represent the median.

### Supplemental method for estimating the founding population in each organ

Abundance of the signature tagged bacteria was determined by using qPCR and CFU/organ data. For each signature tag in each organ, the amount of DNA in ng was calculated by a standard curve using the C_p_ values and known ng amounts. The frequency of abundance of signature tags was determined by dividing the calculated ng of DNA for each tag over the summed total ng for all signature tags in a given organ. The frequencies were then multiplied by the amount of CFU/organ at the time of dissection. Finally, the harmonic mean was calculated for each organ to find the effective population.

